# Spatial dissection of ADC/RPT targets defines therapeutic opportunities in rhabdoid tumors

**DOI:** 10.64898/2025.12.11.692900

**Authors:** Nic G. Reitsam, Victoria Fincke, Maria Daniela Hernandez Ramirez, Marlena Mucha, Eva Sipos, Lisa Siebenhüter, Johanna S. Enke, Maurice Lossner, Christian Vokuhl, Constantin Lapa, Martin Hasselblatt, Michael Frühwald, Bruno Märkl, Pascal Johann

## Abstract

Rhabdoid tumors (RT) are among the most aggressive pediatric malignancies, characterized by early onset, loss of SWI/SNF complex members (SMARCB1 or SMARCA4), and dismal outcomes despite multimodal therapy. Refractory and relapsing RT remain almost uniformly fatal, and targeted or immune-based approaches have yet to demonstrate clinical benefit. To explore novel therapeutic vulnerabilities, we systematically investigated the expression of clinically actionable surface proteins that could serve as targets for antibody–drug conjugates (ADCs), radiopharmaceutical therapy (RPT), or cellular immunotherapies. Based on large-scale transcriptomic analyses, we prioritized FAP, CXCR4, and IL13RA2 and performed comprehensive protein-level validation by immunohistochemistry in an unprecedented cohort of 60 rhabdoid tumors spanning all molecular subgroups (ATRT-TYR, ATRT-SHH, ATRT-MYC, and eMRT). Integrating these data with spatial and single-nucleus transcriptomic profiling, we identified subgroup- and cell-type–specific expression patterns, including heterogeneous FAP distribution between stromal and tumor compartments and a distinct IL13RA2-positive rhabdoid cell population with melanosomal and stem-like features. These findings define a set of biologically and clinically relevant surface targets in RT and provide a translational blueprint for rational ADC and RPT target discovery in pediatric cancer.

## Introduction

Irrespective of their anatomical localization, rhabdoid tumors (RT) are characterised by a dismal prognosis and remain among the most lethal pediatric malignancies despite intensive multimodal therapy. These tumors can occur intracranially (termed atypical teratoid rhabdoid tumors, ATRTs), or extracranially (extracranial malignant rhabdoid tumor, eMRTs). RT usually affect very young children, with a median onset of disease of ca. 1.5 years. The hallmark genetic aberration of these tumors is the loss of SMARCB1 - an integral component of the SWI/SNF complex - which occurs in approximately 98% of all cases (1). The remaining cases are characterized by the loss of SMARCA4. Both alterations are presumed to ultimately inactivate tumor suppressor genes thereby initiating tumorigenesis.

Current therapy combines surgery, intensive chemotherapy, and often radiotherapy, yet outcomes remain poor. Despite multimodal therapy, 60% of children still succumb to their disease, underscoring the urgent need for mechanistically guided and targeted therapies (2). As RTs arise due to a defective chromatin remodelling complex and thus display tractable epigenetic landscape alterations, substantial efforts have been directed towards identifying drugs that may reverse some of the epigenetic changes. A salient example for such a drug is tazemetostat (3)(4)(5), which inhibits the protein EZH2 and may thus limit the oncogenic effect of this enzyme by reactivating tumor suppressor genes. However, due to its limited successes in clinical trials (particularly in SMARCB1-deficient chordomas and sinonasal carcinomas (6)(7)(8), EZH2 inhibitor so far has not proven to be an integral addition in clinical routine therapy, and it is usually only used in relapse settings.

The same is true for tyrosine kinase inhibitors which have emerged from CRISPR-Cas 9 screenings in rhabdoid tumor cell lines and which do not display a relevant activity in the clinical setting (9). Emerging CAR-T strategies targeting ATRT surface antigens have demonstrated preclinical efficacy (10) but none of these approaches have yet translated into clinical application.

Overall, successful attempts to target surface structures in rhabdoid tumors are rare and in contrast to many adult tumor entities such as e.g. bladder cancer (Enfortumab-Vedotin trials as reference(11), breast cancer (TROP2, Sacitumab-Govitecan, (12)), colorectal (13). However, despite these successes in adult oncology, none of these surface targeting modalities, such as antibody-drug conjugates (ADC) or radiopharmaceutical therapy (RPT), have been translated into routine treatment for pediatric tumors, and even compassionate-use access remains extremely limited due to both biological and regulatory constraints.

The broad landscape of clinically actionable ADC and RPT targets (e.g. TROP2, HER2, CXCR4, FAP, etc.) remains virtually unexplored in pediatric oncology. Therefore, in this study, we performed a large-scale investigation of gene expression data in RT to identify potential targets, and identified three genes, whose expression was increased in RT compared to healthy tissues (FAP, CXCR4 and IL13RA2).

FAP is a serine protease, commonly expressed in those fibroblasts that undergo active remodelling processes, such as in fibrosis, autoimmune processes but also in cancer. Its role is best studied in the context of cancer-associated fibroblasts (CAFs) in adult carcinomas with extensive desmoplastic stroma such as pancreatic cancer and colorectal cancer (14). Expressed by activated fibroblasts, it activates secretion of soluble factors such as TGF-b and VEGFA thereby inhibiting dendritic cells and contributing to epithelial to mesenchymal transition (EMT). Its role as a diagnostic marker in nuclear medicine is well established: FAP-directed PET/CT has a high sensitivity in detecting numerous malignancies (15). Beyond this, there is also a therapeutic application by employing radiolabelled therapeutic vectors, e.g. [^177^Lu]Lu-FAP-2286 or ^90^Y-FAPI-46, which have already been shown to be effective in several cancer entities (16,17)(18). Similarly, ADC constructs against the protein are under investigation in Phase I trials, such as e.g. OMTX705 which targets FAP with a tubulysin payload.

Despite extensive data in adult carcinomas, FAP expression patterns and functional roles in pediatric tumors remain largely unknown. The most comprehensive assessment of FAP in sarcomas - although not specifically pediatric tumors - was presented by *Crane et al.* (19) showing high levels of FAP in e.g. osteosarcoma.

More recent literature has shown that MPNSTs express high levels of FAP (*Reitsam et al. in revision)*. It is noteworthy that in some tumor entities FAP is not only expressed by CAFs, but also by the tumor cells themselves. This is particularly observed in sarcomas (20) but also to some degree in mesotheliomas, gastric cancer and other tumor types. Recently, increased FAP expression was linked to the presence of EMT in clear cell renal carcinoma (ccRCC) - thus linked to a particularly aggressive tumor subtype, which points at a potential biological role of increased FAP expression in tumor cells

Like FAP, CXCR4 offers dual diagnostic and therapeutic potential (21), yet its relevance in rhabdoid tumors has not been systematically characterized. The CXCR4 inhibitor Mavorixafor has been employed in clinical studies and several ADC constructs are under evaluation (22). In the rhabdoid tumor context its expression has been detected in cell lines, without a clearly ascribed functional role so far (23). The work which has been conducted in various entities mainly bears functional implications and highlights CXCR4 as a marker for metastatic potential and cancer stem cells (24). IL13RA2 a target that has been investigated in the context of adult glioblastomas (25), adrenocortical carcinomas and melanomas and holds promise of using off-the-shelve CAR-T cells against it. Additionally, IL13RA2 ADC constructs are in active development (26). The physiological function of this which displays in non-neoplastic tissues particularly high expression in testes and ovarian tissue remains not fully elucidated. IL13RA2 has been considered as a decoy receptor for IL13. However, more recent literature points to a signaling activity of the receptor. Moreover, IL13RA2 seems to be associated with a worse prognosis in several different tumor entities (doi: 10.18632/oncotarget.10297, https://doi.org/10.1016/j.bbcan.2022.188802).

In this study, we aimed to assess the viability of FAP, CXCR4, and IL13RA2 as therapeutic targets in rhabdoid tumors. Using immunohistochemistry, we examined the levels of these three proteins in 60 cases, representing ATRT-TYR, ATRT-SHH, ATRT-MYC and eMRT (5) and found that there are subgroup-specific differences. Moreover, FAP protein levels displayed heterogeneous patterns when comparing tumor cells and stroma. Additionally, we leveraged Visium HD spatial transcriptomic data to identify IL13RA2 as a marker for a distinct subset of rhabdoid cells that express melanosomal markers and stem cell markers irrespective of their subgroup and may thus represent a therapeutic target with a peculiar biological role as a tumor driver. Our study provides a translational blueprint for rational ADC and RPT target discovery in pediatric malignancies, bridging tissue-based molecular profiling with therapeutic feasibility.

## Results

### Analysis strategy and approach

To identify therapeutically targetable antigens in rhabdoid tumors, we analysed published expression data from ATRTs and eMRTs (Figure 1A). We examined a curated panel of candidate therapeutic targets (Supplementary Figure 1) and compared their expression in ATRT and eMRT against normal tissue controls from Roth et al. Most antigens associated with endo- or exocrine functions (e.g., CCKBR, SSTR2) showed minimal or absent expression. However, three targets exhibited upregulation in at least one of the ATRT subgroups or eMRT: FAP displayed variable expression across all ATRT subgroups (Figure 1B), CXCR4 demonstrated high expression across all three ATRT molecular subgroups (Figure 1C), and IL13RA2 was significantly upregulated in ATRT-TYR compared to controls (Figure 1D).

**Figure 1.**
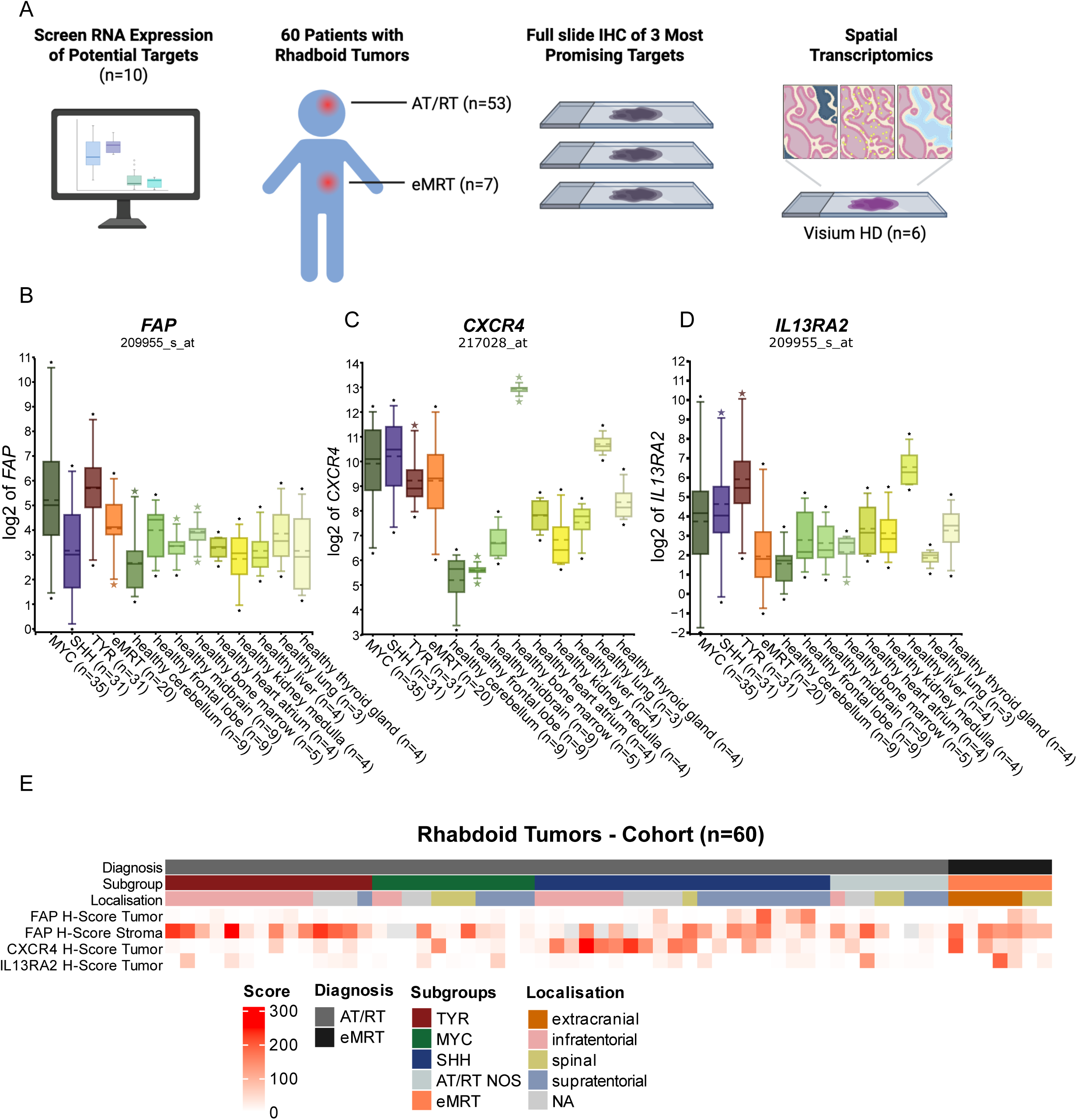
Cohort overview and selection strategy A) Analysis strategy and prioritization of the three target molecules investigated. B-D) Boxplots display the expression levels of the three respective genes as measured by Affymetrix U133 arrays. Leftmost boxes display the three ATRT subgroups, boxplots in yellow and green display normal controls of various tissues E) Overview of the dataset, including molecular subgroup, localization and immunohistochemistry scores.

Based on these results, we focused our next steps on these three proteins and analysed their expression and protein levels in a large number of RT samples (Supplementary Table 1). Overall, the study cohort for full slide immunohistochemical analysis comprised 14 ATRT-TYR, 20 ATRT-SHH, 11 ATRT-MYC, 8 ATRT (unknown subgroup) and 7 eMRT (Figure 1E). Distribution of localisation between the subgroups was found to be as previously described (27).

### Fibroblast activation protein (FAP) shows differential expression in stroma and tumor cells in ATRT-TYR and ATRT-SHH

FAP immunohistochemistry revealed two distinct staining patterns: First, a stromal-predominant pattern, which is typical for malignant epithelial neoplasms with high FAP expression in the cancer-associated fibroblasts (28), was present in 51/60 cases (FAP stromal score > 10), often accentuating perivascular stroma with mostly completely negative tumor cells (Figure 2A). Notably, a strong stromal FAP staining (IHC score >100) was observed in 22 cases. Although distribution of positive cases did not differ significantly between the subgroups (Chi-square test p-value = 0.594), staining in ATRT-TYR cases was stronger than in the other groups (mean H score = 131.46, Figure 2B). Second, we observed FAP expression in the ATRT tumor cells in a subset of cases (16/60 with FAP tumor H-score ≥10, Figure 2 C,D); tumor-cell intrinsic FAP expression is mainly known from mesenchymal neoplasms *(Crane et al., Reitsam et al. in revision)*. This expression pattern was characterized by a focal but strong cytoplasmic and membranous FAP expression. Overall, there was no significant correlation between tumors that displayed stromal and tumor FAP positivity (Figure 1 E). In the corresponding cases, tumor cell-specific FAP expression was always only detectable in a subpopulation of tumor cells and never in the entire tumor (maximum FAP H-scores: 190). Notably, within these tumors, rhabdoid cells were preferentially highlighted by FAP. Similar to the stromal positive cases, there was a predominance of ATRT-SHH (7 cases) and the two cases with a strikingly positive staining for the tumor belonged to ATRT-SHH.

**Figure 2:**
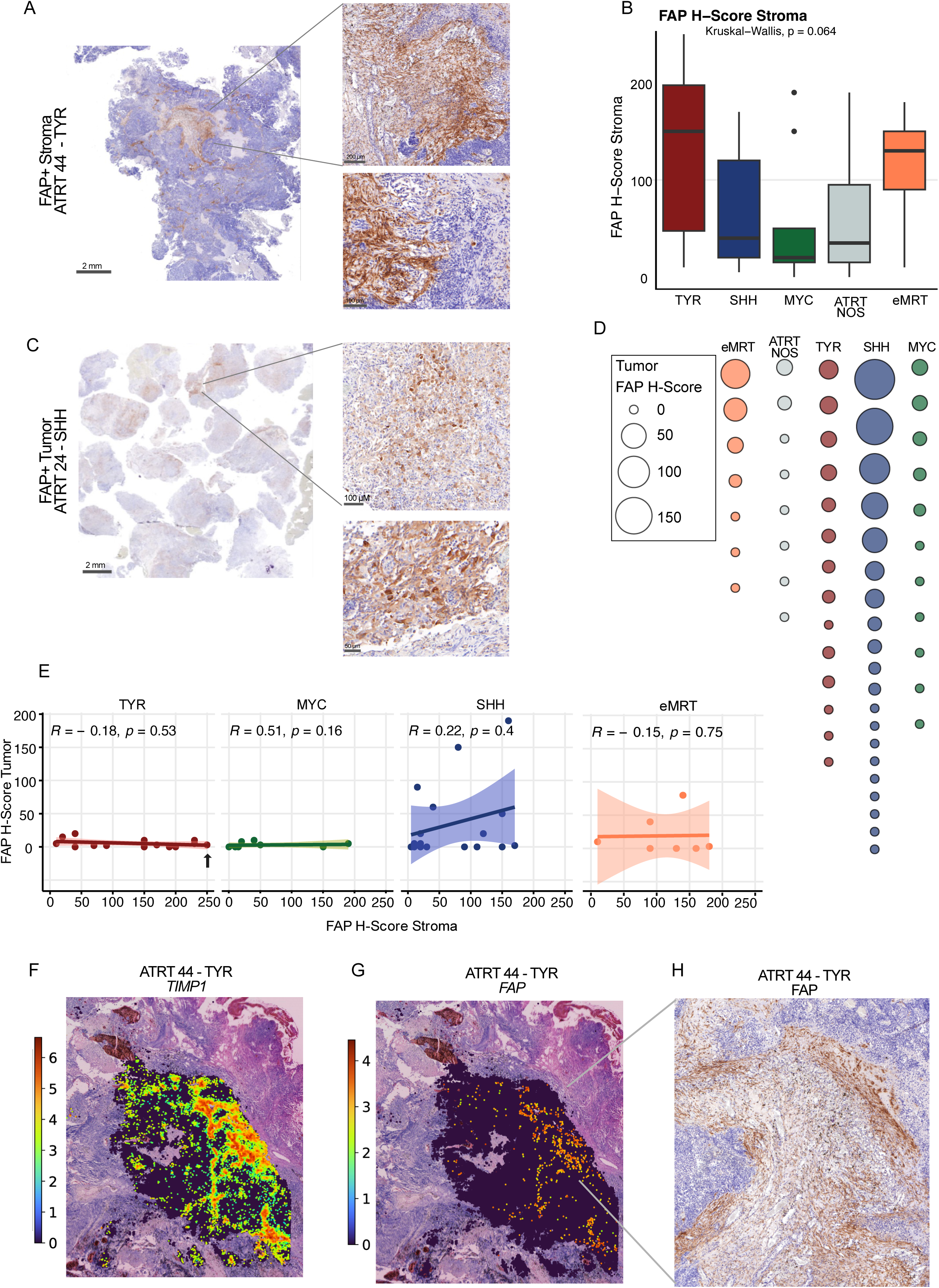
FAP IHC staining patterns in rhabdoid tumors A) Example for a tumor displaying strong staining pattern of FAP in the tumor stroma B) Box plots displaying FAP H-Scores for stroma compartment by subgroup C) Example for a tumor displaying FAP staining in tumor cells D) Bubble plot demonstrating the predominant expression of FAP ATRT-SHH samples D) Line plots show the correlation between FAP tumor cell positivity and stromal positivity in the respective subgroups E) TIMP-1 expression correlates with FAP expression in areas of stromal infiltration F) and G) Images display areas of FAP positivity in spatial transcriptomics (F) and in immunohistochemistry (G) H) IHC FAP staining corresponding to the same are where *FAP* expression is observed

Aiming to resolve FAP expression on cell type level, we performed spatial correlation analyses of FAP positive stromal cells using the Visium HD system. In accordance with the immunohistochemical stainings, we detected focal FAP positivity often colocalising with TIMP1 (Tissue Inhibitor of Melloproteinases 1; Figure 2 F,G), which is known to be upregulated in cancer-associated fibroblasts (29) and known to induce aberrant blood-vessel formation and anti-apoptotic activity. RNA expression of FAP was validated by protein expression in the same are (Figure 2H).

Next, we analysed published ATRT single nucleus RNA-Seq dataset (*30*), we could validated our protein findings proving that FAP expression is heterogeneous between ATRT tumor samples (Supplementary Figure 4) and found expression mostly in the stromal compartment of single patients.

The second type of FAP expression - positivity in the tumor cells - was not detected in our spatially resolved cohort, most likely owing to the low incidence in the IHC cohort.

### CXCR4 expression is detected in tumor cells and tumor vasculature

Across the cohort, a relevant subset of 34 cases (CXCR4 tumor H-score ≥10) displayed a focal but strong membranous CXCR4 expression on the tumor cells (Figure 3A). Among these were 16 cases belonged to ATRT-SHH subgroup. Again, ATRT-MYC was underrepresented (only six cases stained positive, with comparably low IHC scores between 10-20). Only one case of ATRT-MYC (localized to the spine) showed a higher CXCR4 expression on the tumor cells (H-score: 140, Figure 3B). By comparing semiquantitative, H-scores, we could validate that CXCR4 tumor cell positivity was higher in ATRT-SHH than in the other ATRT subgroups (p<0.05, respectively) (Supplementary Table 2, Supplementary Figure 2).

**Figure 3:**
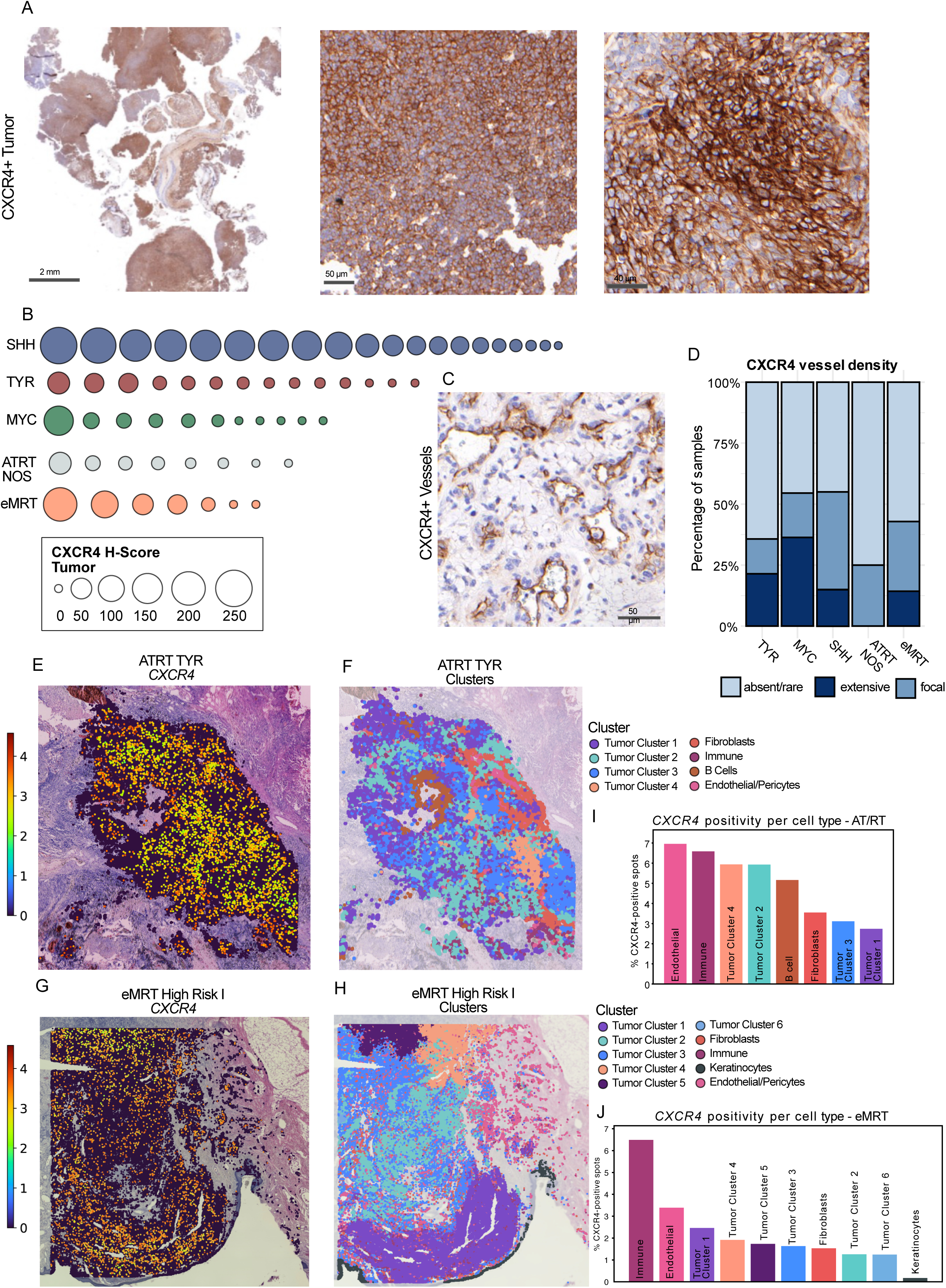
CXCR4 staining patterns in rhabdoid tumors A) Example of a tumor displaying strong staining pattern of the tumor cells B) Bubble plot demonstrating the protein expression of CXCR4 in ATRT-SHH C) Tumor with a preferentially perivascular staining of CXCR4 D) Staining pattern of blood vessels for CXCR4 is depicted in a stacked barplot and graded into “extensive”, “absent” and “focal” E-F) *CXCR4* expression in an ATRT-TYR and the decomposition into clusters G-H) *CXCR4* expression in an eMRT High Risk I and the decomposition into clusters H-I) Barplots showing the CXCR4 positivity by cell type for an ATRT TYR and an eMRT High Risk I

Interestingly, 11 cases showed a high CXCR4-positive vessel density characterized by a strong CXCR4 expression in endothelial cells of a dense capillary network between the tumor cells (Figure 3 C), which was already described for HCCs by *Xu et al.* and was shown to have a predictive value for antiangiogenic treatment such as sorafenib (31). Notably, CXCR4-positive vessel density seems to display a balanced pattern across the subgroups (3 cases ATRT-TYR, 3 cases ATRT-SHH, 3 eMRT, 1 ATRT-MYC, chi square test n.s.)(Figure 3D, Supplementary Table 3).

The widespread expression of CXCR4 on the protein level in the cohort prompted us to further investigate the cell populations that may be targeted by CXCR4-directed therapies with spatial resolution. Spatially resolved single-cell transcriptomic analysis in one ATRT-TYR (Figure 3 E,F) and one eMRT (Figure 3 G,H) revealed pronounced *CXCR4* expression across multiple cellular compartments. The highest proportions of *CXCR4*-positive cells were detected in endothelial cells, followed by immune cells, tumor cluster 4 (corresponding to an immune-reactive phenotype), tumor cluster 2 (characterized by invasive-like features), and B cells (Figure 3 I-J). This spatially distinct expression pattern was recapitulated in eMRT tissue samples, where *CXCR4* expression was most prominent in immune cells, endothelial cells, tumor clusters with mesenchymal/developmental-like features, and clusters with immune-modulating phenotypes. Protein level expression mirrored our spatial transcriptomic findings with CXCR4 expression not only in tumor cells but also in intermingled immune cells and, as described, in some cases in tumor-associated vessels.

Spatial feature plots and quantitative analyses in one eMRT and one ATRT-TYR demonstrated differences in *CXCR4* and *CXCL12* (ligand of CXCR4) gene expression across annotated tissue hotspots: In hotspot 1, high proportions of *CXCL12* and *CXCR4* co-expression were observed in both tumor and endothelial cell populations, suggesting local, paracrine ligand-receptor interaction (Figure 4 A, B, E). In contrast, hotspot 2 showed elevated *CXCL12* expression in stromal and vascular compartments with only *CXCR4* expressing tumor cluster 4 cells, supporting a primarily paracrine signaling model (Figure 4 C, D, F).

**Figure 4.**
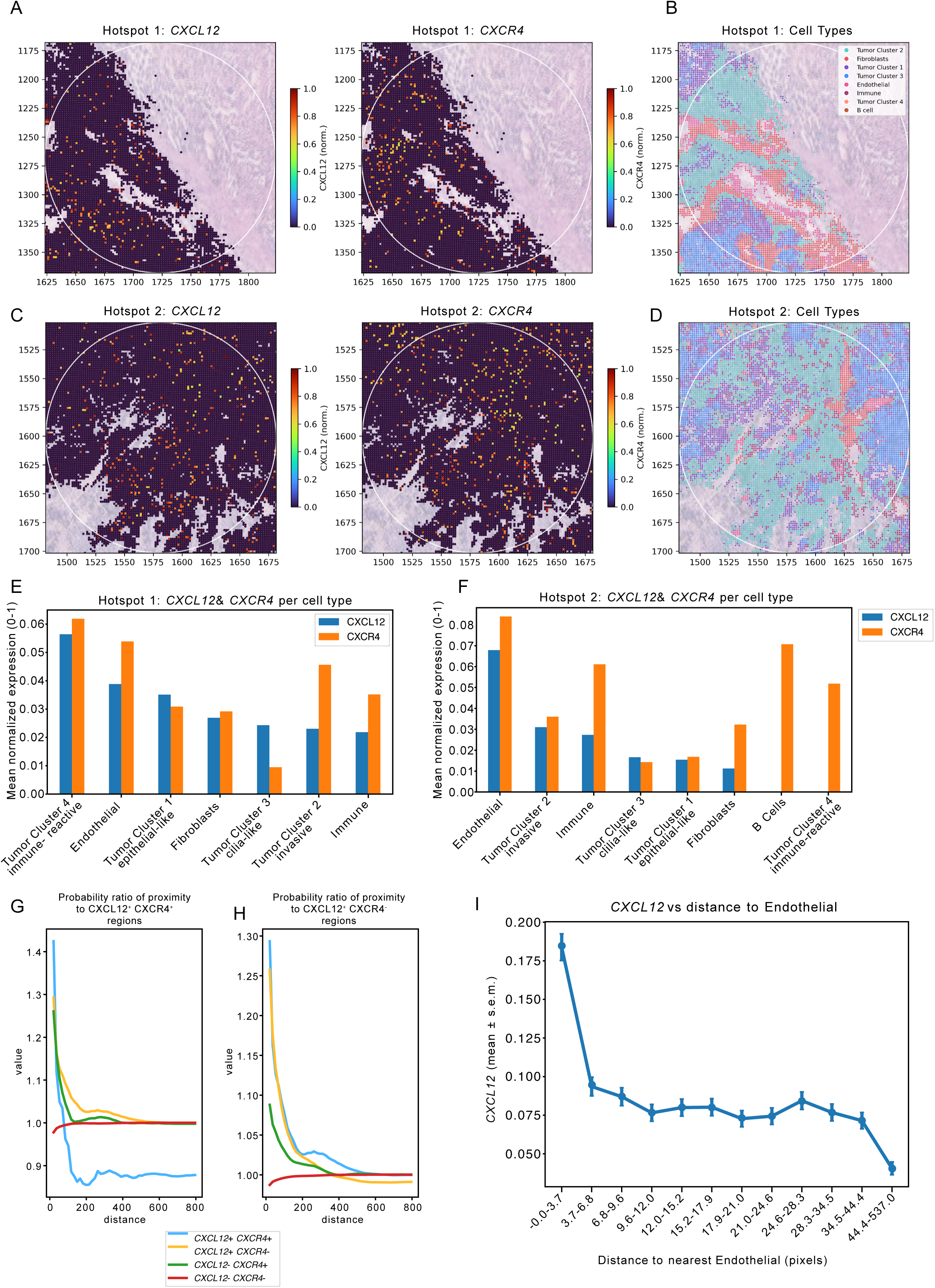
Spatial *CXCL12-CXCR4* co-localization analysis in rhabdoid tumors. A-D) Spatial distribution of *CXCL12* (A, C) and *CXCR4* (B, D) expression with cell typeannotations in two tumor hotspots overlaid on H&E. Hotspot 1 shows ligand-receptor co-expression in tumor and endothelial cells; Hotspot 2 displays CXCL12 enrichment in stroma/vasculature with diOerential CXCR4 patterns. E, F) Mean normalized *CXCL12* and *CXCR4* expression across cell types in Hotspot 1 (E) and Hotspot 2 (F) G, H) Spatial proximity analysis showing probability ratios of co-localization between *CXCL12/CXCR4* expression states. *CXCL12+CXCR4+* cells exhibit highest spatial association, indicating active ligand-receptor pairing. I) CXCL12 expression versus distance to endothelial cells. CXCL12 levels peak near vasculature, identifying endothelial cells as a major ligand source. Data: mean ± s.e.m.

Co-localisation analyses revealed that *CXCL12*+ and *CXCR4*+ cells displayed the highest degree of spatial proximity (Figure 4G), compared to *CXCL12* and *CXCR4* double negative and single negative cells respectively. Cells expressing either *CXCL12* or *CXCR4* alone were also more likely to localize near other ligand-expressing cells, while *CXCL12*+*CXCR4*-cells displayed only limited co-localization with *CXCL12*-*CXCR4*+ populations, but showed notably greater colocalization with any *CXCL12*+ cells (Figure 4H). Further analysis demonstrated a clear gradient of *CXCL12* expression, with levels decreasing as the distance from endothelial cells increased. This spatial organization suggests that endothelial cells are the principal *CXCL12* producers within ATRT tissue, creating chemokine-rich niches that facilitate the attraction or retention of *CXCR4*-expressing tumor and immune cells (Figure 4 I).

Overall, unlike FAP with a more focal pattern, the widespread expression of CXCR4 holds the promise to target not only compartments of the tumor but both tumor-associated blood vessels (as a putative contributor to cancer spread) and tumor itself.

### IL13RA2 Marks a Neural-Crest–Like, Stem-Associated Subpopulation in ATRTs

IL13RA2 has been explored extensively in adult gliomas as a potential target for CAR-T cell therapy. The potential for a rapid translation into the clinic further reinforces its attractiveness as a target. In total, 11 samples stained positive for IL13RA2 when applying the previously published threshold (H-score ≥20 as inclusion criteria for CAR-T trial in gliomas) (Figure 5 A,B). Unlike in the two other target molecules, we found the positive cases to be equally distributed over the tumor groups (Figure 5 C).

**Figure 5.**
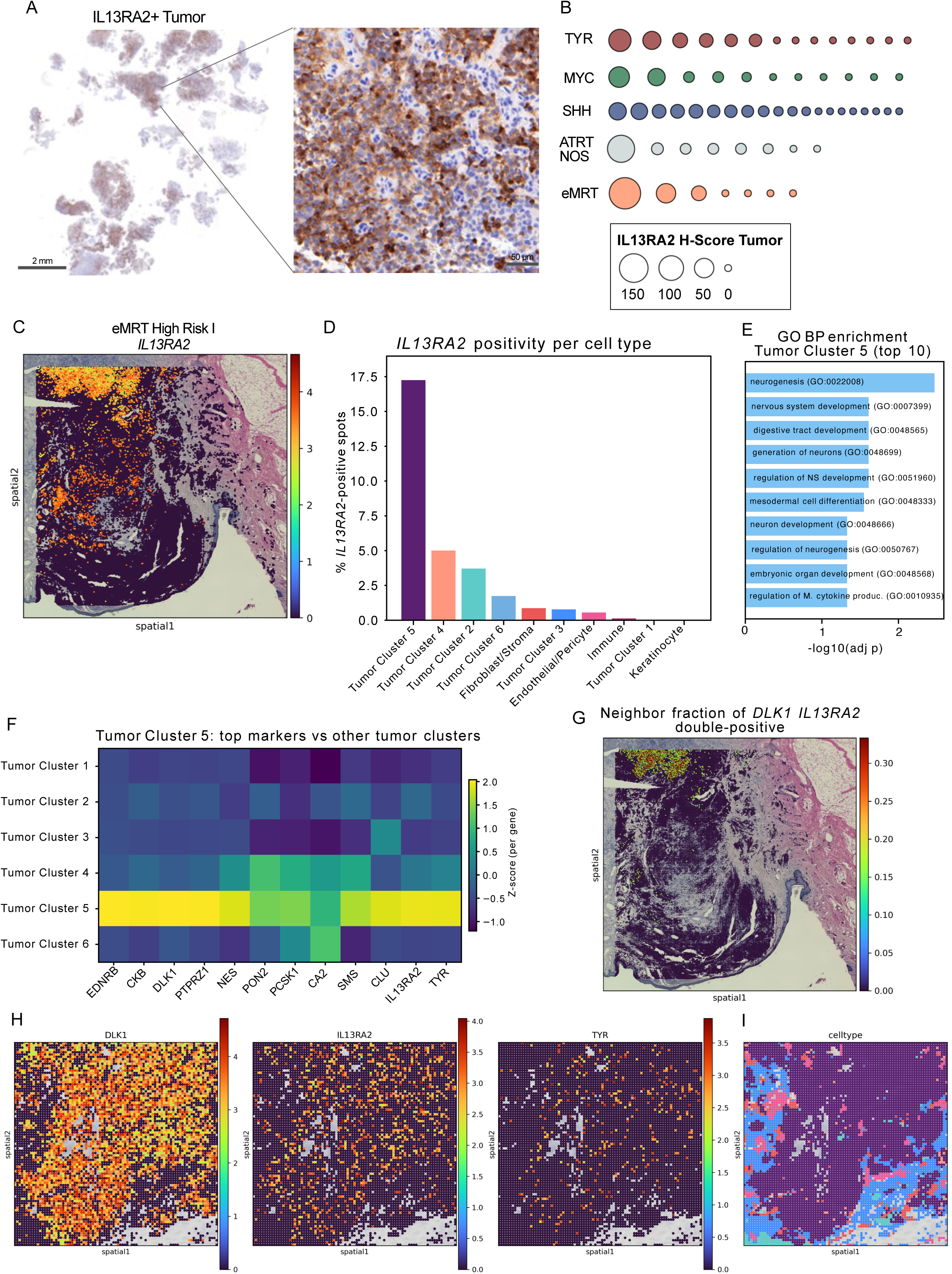
Immunohistochemistry and intra sample distribution of IL13RA2 A) Immunohistochemistry of Il13RA2 displaying positive tumor cells in one sample B) Bubble plot displaying Il13RA2 IHC score per subgroup and sample C) Spatial mapping of IL13RA2 expression in an eMRT sample D) Il13RA2 expression is present particularly in one tumor cluster E) GO term enrichment analysis of IL13RA2 expressing tumor cluster F) Genes that are coexpressed with IL13RA2 G) Coexpression of DLK1 and IL13RA2 H-I) Close up of the spatial distribution of *DLK1, IL13RA2* and *TYR* expression and display of tumor cluster

When investigating the distribution of IL13RA2 spatially, we found it to be confined to subclusters of the tumor cells: In the eMRT case investigated, *IL13RA2* expression marked a specific cluster of tumor-cells (Figure 5 D, E) with neural crest-like signatures (Figure 5 F) characterised by a high expression of melanosomal markers (such as TYR and MITF), but also stem cell markers such as nestin (Figure 5 G). Interestingly, this cell population displayed a high expression of *DLK1* (Figure 5 H), a member of the NOTCH signaling pathway which has been recently identified as an immunotherapeutic target in other entities (32). While IL13RA2 expressing cells are also present in Tumor Cluster 4 and 2, the co-expression with DLK1 is strictly limited to Tumor cluster 5 (Figure 5 G-H). While cluster 5 tumor cells almost uniformly express DLK1, IL13RA2 and TYR positive cells are mixed into the cluster, pointing at a certain degree of heterogeneity even within tumor cells (Figure 5I, J).

The pattern of TYR and IL13RA2 coexpression was also observed in a case of ATRT (subgroup TYR, Figure 6 A). Here, the presence of a *TYR* positive cell population is less surprising, given the previously described enrichment in melanosomal markers (33)(27)(34). We found the *IL13RA2/TYR* positive population to be highly enriched in proliferation associated terms, but also in gene sets that associate with cilial assembly (30) when compared to only *TYR* expressing cells (Figure 6 B). *IL13RA2/TYR* coexpression was found again almost exclusively in the tumor (Figure 6 C, D) and not in the stromal compartment (Figure 6 E). Investigating the ligand receptor interactions between *IL13RA2* and *TYR* co-expressing cells, revealed several activated pathways. The TYR GRP143 axis is active in signalling to all surrounding cell types, while MIF to CD74 is limited to fibroblasts and immune cells, and IFNG to CD74 to immune cells only (Figure 6 F). Expression of CD74 is spatially elevated in Fibroblast and Immune cell rich parts (Figure 6 G), neighboring cells with IL13RA2 and TYR expression (Figure 6 H)

**Figure 6.**
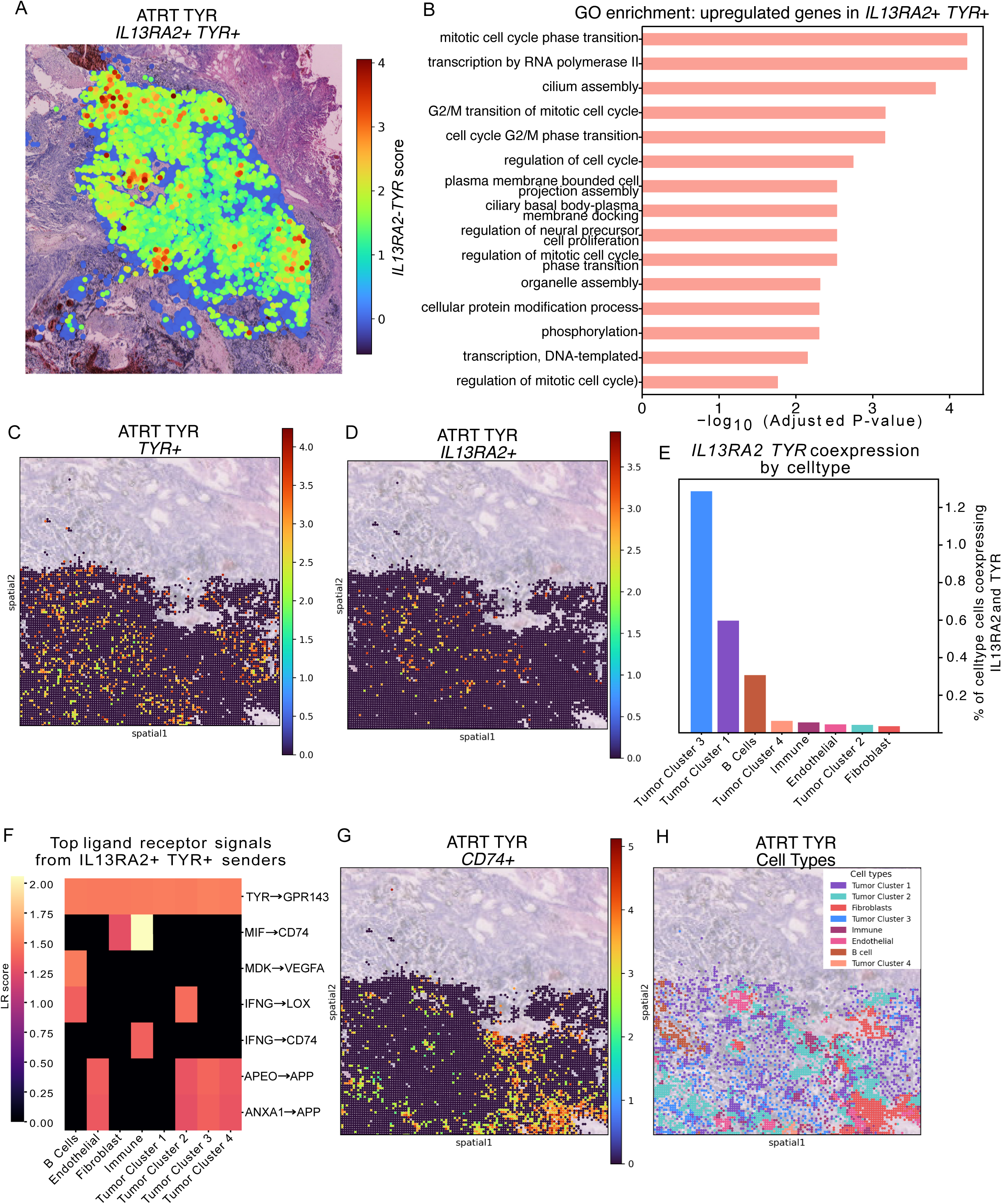
Characterization of IL13RA2 expressing cells in an ATRT-TYR sample A) Colocalization of IL13RA2 and TYR in an ATRT TYR sample B) Gene set enrichment analysis of TYR and IL13RA2 double-positive cells C-D) Spatial mapping of *IL13RA2* and *TYR* expression in an ATRT-TYR sample E) Distribution of IL13RA2 and TYR co-expression over the cell types F) Ligand-receptor signaling in IL13RA2 and TYR positive cells G-H) Spatial expression of CD74 and cell type distribution

Overall, our findings suggest heterogeneous target expression of the aforementioned antigens and thus reinforce the need for thorough immunohistochemical analysis in each case that qualifies for RPT or ADC therapy.

## Discussion

While the histopathological diagnosis of rhabdoid tumors is usually unambiguous, the further course of disease is often marked by uncertainty: particularly in cases with early progression, which occur frequently, it remains unclear which salvage therapies are most effective. Given the limited success of epigenetically acting agents such as DNMTi (35) or tazemetostat (36) - the latter showing only few partial responses in rhabdoid tumors - the best treatment modality in relapsed or refractory cases remains unclear.

Inspired by successes in adult cancers, the last decade has seen ADC constructs gradually gaining ground in pediatric oncology. The most prominent example certainly being anti-GD2 directed antibody therapies that are part of routine therapy for neuroblastomas. Increasingly, the antibody Dinutuximab is also used in relapse settings of other solid tumors including Ewing sarcoma or osteosarcoma, but so far without clear evidence for efficacy (37). Another major target which has been under active investigation is B7-H3, also in rhabdoid tumors where preclinical studies have motivated the use of CAR-T cells in the CSF to target tumor cells (10).

Our approach based on previously published gene expression data, suggests an upregulation of FAP, CXCR4 and IL13RA2 in rhabdoid tumors. Based on this, our current study provides a systematic evaluation of these three clinically actionable targets in a large, well-characterized cohort (n=60) of pediatric RTs combined with snRNA and spatial transcriptomics data. Employing whole-slide IHC rather than TMAs minimized sampling bias and enabled high-resolution quantification of sub-clonal tumor-cell expression patterns and stromal architecture - including perivascular niches - thus improving biological interpretability (38) and correlation to other -omics modalities.

The identification of FAP as a relevant target in ATRT is particularly unexpected, as FAP is typically characteristic of desmoplastic, stroma-rich malignancies such as pancreatic adenocarcinoma (39) or colorectal carcinomas (40), and not brain tumors. Based on our spatial and snRNA data, we have shown that in a subset of RTs FAP is mainly expressed in cancer associated fibroblasts.

At the protein level, approximately half of all RT patients display an expression either of FAP in the stroma or on the tumor cells. The exact threshold which is to be used when it comes to directing therapies against this target in clinical trials is still unclear; however our study motivates further investigations - the possibility to use Lutetium-177 or Actinium-225 coupled FAP directed antibodies in progressive disease, characterised by a disruption of the blood brain barrier is thus apparent (41). Whether the addition of a “stroma-directed” therapy component may be advisable, needs to be tested in clinical trials combining this modality and classical chemotherapeutic approaches - stroma-targeting would display a completely novel treatment paradigm in ATRTs, potentially eradicating another disease-driving mechanism. Future work should clarify whether tumor-cell versus stromal FAP expression conveys distinct biological or therapeutic implications. For other tumor entities, it is known that stromal FAP expression conveys immunosuppressive properties (42) while tumor-cell intrinsic FAP expression seems to be linked to an EMT-like phenotype ((43), *Reitsam et al*., in revision).

In contrast to FAP, CXCR4 was broadly expressed across our ATRT cohort, albeit with substantial inter- and intratumoral heterogeneity. By our multi-omic approach, we could confidently show that CXCR4 expression in ATRTs spans tumor cells, tumor-associated vasculature and interspersed immune cells. From a theranostic point of view, CXCR4 has not only the potential for molecular whole-body in-vivo imaging (44) but also to be a target for CXCR4-directed endoradiotherapy. Notably, a 11 out of 60 rhabdoid tumors displayed a remarkably high density of CXCR4-positive vessels, characterised by strong endothelial staining within a dense capillary network intercalated between the tumor cells. A similar vascular CXCR4 phenotype has previously been described for hepatocellular carcinoma by (45) where it was functionally linked to poor prognosis, pro-angiogenic signalling and predictive of response to anti-angiogenic therapy with sorafenib. These findings suggest that endothelial CXCR4 expression in ATRTs may likewise reflect a so-far unknown state of active angiogenic signalling. Indeed, early functional studies in ATRT cell lines demonstrated activation of diverse tyrosine kinases, among them VEGF and SDF-1/CXCL12, along with growth inhibition upon sorafenib or sunitinib exposure (*46*). Together with our spatial data showing close colocalization of CXCR4-expressing endothelial cells, these observations imply that CXCR4 contributes to a pro-angiogenic microenvironment in ATRT and may serve as a dual-compartment target, with potential sensitivity to CXCR4-directed or multi-kinase-based anti-angiogenic strategies. From a clinical standpoint, IL13RA2 marks the most mature target, in the sense that it is undergoing extensive clinical evaluation in Glioblastoma, using CAR-T cells as therapeutic modality.

Importantly, while we did not directly assess receptor internalization or internalization kinetics in ATRTs, such functional properties are a prerequisite for both radioligand therapy and many ADCs. However, extensive preclinical work in other tumor types, including CXCR4-directed RPT (47) and FAP-directed approaches (48), has demonstrated robust internalization in CXCR4+ and FAP+ cells, with internalization efficiency scaling with expression levels. Although these data have not been generated specifically in ATRTs, the analogy is biologically well supported and sufficient from a nuclear medicine perspective to justify therapeutic exploration of these targets.

Clinically, IL13RA2 represents the most mature of the three targets, owing to extensive evaluation in glioblastoma using CAR-T cells. In our cohort, IL13RA2 emerged as a tumor-restricted, clonally focal antigen: 11/60 ATRTs would meet the clinically used inclusion threshold (H-score ≥20) used in the already mentioned IL13RA2-CAR-T trial in high-grade gliomas. Spatial and single-cell data linked *IL13RA2* expression closely to a neural-crest like, stem-associated population with *TYR, MITF* and *DLK1* co-expression, suggesting IL13RA2 marks developmentally primitive, proliferative niche than a generic bulk phenotype. Hence, IL13RA2 represents an attractive therapeutic entry point to eradicate proliferative, stem-associated ATRT cells. Noteworthy, besides the described IL13RA2 CAR-T product (49) also IL13RA2-guided ADCs (50) as well as theranostic approaches (51) are under active development.

Recent insights into the molecular and immunological mechanisms of ADCs (52) but also RPT (53) highlight their efficacy extends well beyond direct cytotoxic effects. Those targeted therapies can trigger immunogenic cell death, activate dendritic cells and actively reshape the TME, thereby synergising with immune checkpoint blockade and other immunomodulatory therapies. In the context of our study, which identifies a heterogeneous stromal and tumor-cell expression of the investigated targets in ATRTs, these findings are particularly relevant: as shown, each of these targets resides at intersection of tumor-stroma-immune interactions. Thus, FAP-, CXCR4-, IL13RA2-directed therapies could not only deliver direct toxicity in a subset of ATRTs but also modulate immune tone, offering a biological rationale for combination strategies.

In summary, our study fills a significant gap in the characterisation of novel therapeutic targets in rhabdoid tumors. Given the current scarcity of such analyses in RTs, and with rapid advances in CAR-T cells, ADCs, and theranostic technologies, we anticipate that the coming years will see increased efforts to translate and refine these approaches across pediatric solid tumors.

### Strengths and Limitations

A major strength of this study lies in its comprehensive, multimodal assessment of three clinically actionable surface antigens, FAP, CXCR4 and IL13RA2, across an unprecedentedly large and well-annotated rhabdoid tumor cohort (n=60). By employing whole-slide IHC, we captured intratumoral and stromal heterogeneity with spatial resolution and integrated these findings with single-nucleus and spatial transcriptomic data, establishing a generalizable blueprint for ADC and RPT target discovery in rare pediatric cancers.

Nevertheless, the study has some limitations. First, target preselection from transcriptomic datasets may have overlooked low abundance yet stable surface epitopes. Second, the observational nature of our tissue-based study limits functional inference about downstream signaling or therapeutic susceptibilities. Third, as already briefly mentioned, threshold for therapeutic relevance (e.g., IHC H-score cut-offs) remain to be established in clinical trials. As another limitation, assessments relied on expert visual scoring; although reproducible and reflecting clinical reality, future studies incorporating digital image analysis with quantitative algorithms, especially refined computational pathology biomarkers, in analogy to normalized membrane metrics (NMR) for the ADC-target TROP2 (54), may further enhance precision and potentially predictive performance.

Overall, our study has shown that target expression for the three proteins investigated may be heterogeneous: This refers to a subgroup-wise difference in distribution but also to the intratumorally heterogeneous distribution pattern of the epitopes. A caveat of our work is the preselection of surface target molecules based on gene expression data: While these might be a useful surrogate for prioritization, the approach may fall short in detecting stable, membrane-bound epitopes with a low mRNA yield but a high stability in the cell or at the cell membrane. In our approach, we have balanced the aim to include as many rhabdoid tumor cases as possible versus a higher number of targets and decided for a more comprehensive evaluation of a limited number of molecules. An unresolved question in the field is the cut-off to use in immunohistochemistry for each of the evaluated targets, in terms of trial inclusion and therapeutic response; only for IL13RA2-directed CAR-T therapy in high-grade gliomas a cut-off of H-score ≥20 for trial inclusion was previously defined (55).

## Methods

### Ethical Approval, Cohort & Samples

Material from 60 patients enrolled in the EURHAB registry (IRB approval no. 2009-532-f-S) was included in this study. Patients and/or legal guardians provided written informed consent. The study was conducted in accordance with the Declaration of Helsinki.

Immunohistochemical staining for IL13RA2, CXCR4, and FAP was performed on 2-µm–thick whole-slide FFPE sections. Whole-slide staining was selected to minimise sampling bias due to intratumoral heterogeneity and to preserve spatial context, including tumor–stroma interfaces and perivascular niches (56).

### Immunohistochemistry for FAP, CXCR4 and IL13RA2

Immunohistochemical staining was performed on a Leica Bond RX automated platform using Bond Polymer Refine Detection Kit from Leica. The following primary antibodies were applied: FAP (clone ab207178, Abcam, dilution 1:400), CXCR4 (ab181020, Abcam, 1:2000), and IL13RA2 (clone E7U7B, Cell Signaling, 1:200). Antigen retrieval was carried out in EDTA buffer at 95 °C, followed by primary antibody incubation at room temperature for 32 minutes.

FAP expression was quantified separately in tumor cells and stromal compartments, given its established dual localisation in cancer-associated fibroblasts (CAF) and, in selected tumor entities such as sarcomas, in malignant cells themselves (19). For CXCR4 and IL13RA2, only tumor cell staining was evaluated. All evaluations were performed visually on immunohistochemical whole-slide sections. Scoring was conducted by the co-first author (NGR), a fourth-year pathology resident with substantial experience in translational pathology research as well as diagnostic immunohistochemical assessment. NGR performed all primary evaluations while fully blinded to subgroup allocation and clinical outcome data. In diagnostically challenging or ambiguous cases, slides were jointly reviewed with BM, a board-certified pathologist with more than 20 years of experience in diagnostic pathology (general and neuropathology cases) as well as research, and a consensus decision was reached.

Staining intensity was semi quantitatively assessed using the H-score formula: *(1 × percentage of weak staining) + (2 × percentage of moderate staining) + (3 × percentage of strong staining)*, yielding a score from 0 to 300 (57)

As expected from prior reports CXCR4 immunoreactivity was occasionally observed in vascular endothelial cells and in normal neural tissues such as the cerebellar granule cell layer (58) and ependymal lining (59). For CXCR4, we also assessed the density of CXCR4-positive vessels in a binary (high vs low) and a three-tiered (rare/absent vs focal vs extensive) approach, similar to what was published by (https://doi.org/10.1158/1078-0432.CCR-16-2131) *Jing et al.* . IL13RA2 showed tumor-restricted expression and was consistently negative in adjacent non-neoplastic tissue, including stroma and vessels. We applied an H-score threshold of 20 (corresponding to ≥20% of tumor cells with 1+ staining intensity) to define IL13RA2 positivity, consistent with the cutoff used for patient inclusion in the IL13RA2-targeted CAR-T cell trial in recurrent high-grade glioma, published by *Brown et al*. (60). The used IL-13RA2/CD213a2 (E7U7B) antibody does not show cross-reactivity with IL13RA1, which shows a relevant homology with the RA2 subunit.

### Spatial Transcriptomics Data Acquisition and Processing

Visium spatial gene expression profiling was performed on 6 tissues from rhabdoid tumors. Tissue sections were processed according to the 10x Genomics Visium HD Spatial Gene Expression protocol. Raw sequencing data were processed using Space Ranger (10x Genomics) to generate feature-barcode matrices and align spots to H&E-stained histology images. Spatial coordinates for each spot were extracted at full resolution for downstream overlay analysis.

### Quality Control and Normalization

Gene expression matrices were imported into Python using Scanpy (v1.9+) and quality control filtering was applied. Raw count data were preserved in a separate layer prior to normalization. Gene expression was normalized using standard log-normalization (log1p transformation) for visualization and differential expression analysis. For gene set scoring and correlation analyses, the raw count matrix was retained to ensure accurate quantification.

### Unsupervised Clustering and Cell Type Annotation

Principal component analysis (PCA) was performed on highly variable genes, and a k-nearest neighbors graph was constructed in PCA space using Scanpy. Leiden clustering was applied to identify transcriptionally distinct spatial domains. Clusters were annotated based on marker gene expression patterns.

### Differential Expression Analysis

Differential gene expression analysis was performed using Scanpy’s rank_genes_groups function with the Wilcoxon rank-sum test. For tumor-specific analyses, only spots annotated as tumor clusters were retained. To identify markers of special Tumor Clusters, differential expression was performed comparing clusters versus all other tumor clusters. Significantly differentially expressed genes were defined as those with adjusted p-value < 0.05, log2 fold change > 0.5, and detection in >5% of spots in the group. Top markers were visualized using dot plots showing mean expression and percentage of expressing spots per cluster.

### Gene Set Scoring

Custom gene signatures were compiled for immune cell populations (total immune, T cells, macrophages), fibroblast subtypes, and other cell types based on canonical markers. Gene set scores were calculated using Scanpy’s score_genes function, which computes the average expression of genes in the signature relative to a control gene set. Scores were computed on the full gene matrix to avoid bias from feature selection during preprocessing. Spatial distribution of signature scores was visualized on tissue coordinates.

### Spatial Neighbor Analysis

Spatial neighborhood graphs were constructed using Squidpy’s spatial_neighbors function with Delaunay triangulation (coord_type=“generic”). The resulting spatial connectivities matrix was symmetrized and binarized to define adjacency relationships between spots. For ligand-receptor co-localisationation analyses (e.g., CXCL12-CXCR4), spots were classified as ligand-positive or receptor-positive based on expression thresholds. Spatial co-occurrence was quantified by counting the number of ligand-positive spots adjacent to receptor-positive spots and vice versa using the spatial graph.

### Gene Co-expression Analysis

To identify genes co-expressed with target genes (e.g., *CXCR4*), Pearson correlation coefficients were calculated between the target gene and all other genes across tumor spots. Genes with minimum detection in ≥50 spots were retained. The top N positively correlated genes (typically top 10, correlation > 0.3) were selected and visualized using heatmaps and dot plots.

### Spatial Gene Expression Visualization

Gene expression was visualized on tissue coordinates using Scanpy’s spatial plotting functions (sc.pl.spatial). Spots were colored by expression level using continuous color maps with size adjusted for high-definition visualization (spot_size typically 6-12 depending on image resolution). Multi-gene panels were generated showing expression of key markers (e.g., *IL13RA2, CXCR4, FAP, CXCL12, DLK1*) overlaid on H&E histology images at high-resolution.

### Tumor Hotspot Identification

For identification of gene expression hotspots (e.g., DLK1/IL13RA2 double-positive regions), individual gene expression values were normalized to z-scores, and a combined score was computed as the sum of z-scores. Spots in the top percentile (e.g., 97.5th percentile) were classified as hotspot spots. Spatial subsets were extracted by either defining a bounding box around hotspot spots with optional padding (bbox mode) or by selecting all spots within a defined radius (e.g., 200 pixels) of hotspot centroids (radius mode).

### Statistical Analysis and Visualization

Statistical analyses were performed using Python scientific computing libraries including NumPy, SciPy, and pandas. Visualization was performed using Matplotlib and Seaborn with custom color palettes applied consistently across celltypes. Figures were exported in both PNG and PDF formats for publication. Dot plots, heatmaps, and violin plots were generated to summarize cluster-specific gene expression patterns.

### Software and Computational Environment

All analyses were performed in Python 3.8+ using Jupyter notebooks. Key packages included Scanpy (v1.9+), Squidpy (v1.2+), NumPy (v1.21+), pandas (v1.3+), SciPy (v1.7+), Matplotlib (v3.4+), and Seaborn (v0.11+). Spatial transcriptomics data were managed using the AnnData object structure.

